# A cellular and molecular portrait of endometriosis subtypes

**DOI:** 10.1101/2021.05.20.445037

**Authors:** Marcos A.S. Fonseca, Kelly N. Wright, Xianzhi Lin, Forough Abbasi, Marcela Haro, Jennifer Sun, Lourdes Hernandez, Natasha L. Orr, Jooyoon Hong, Yunhee Choi-Kuaea, Horacio M. Maluf, Bonnie L. Balzer, Ilana Cass, Mireille Truong, Yemin Wang, Margareta D. Pisarska, Huy Dinh, Amal EL-Naggar, David Huntsman, Michael S. Anglesio, Marc T. Goodman, Fabiola Medeiros, Matthew Siedhoff, Kate Lawrenson

**Author notes:** Correspondence to: Kate Lawrenson, PhD. 290W 3. equal contribution (Marcos A.S. Fonseca, Kelly N. Wright, Xianzhi Lin). jointly directed the study (Fabiola Medeiros, Matthew Siedhoff, Kate Lawrenson).

## Abstract

Endometriosis is a common, benign condition characterized by extensive heterogeneity in lesion appearance and patient symptoms. We profiled transcriptomes of 207,949 individual cells from endometriomata (n=7), extra-ovarian endometriosis (n=19), eutopic endometrium (n=4), unaffected ovary (n=1) and endometriosis-free peritoneum (n=4) to create a cellular atlas of endometrial-type epithelial cells, endometrial-type stromal cells and microenvironmental cell populations across tissue sites. Signatures of endometrial-type epithelium and stroma differed markedly across eutopic endometrium, endometrioma, superficial extra-ovarian disease and deep infiltrating endometriosis, suggesting that extensive transcriptional reprogramming is a core component of the disease process. Endometriomas were notable for the dysregulation of pro-inflammatory pathways and upregulation of complement proteins *C3* and *C7*. Somatic *ARID1A* mutation in epithelial cells was associated with upregulation of pro-angiogenic factor *SOX17* and remodeling of the endothelial cell compartment. Finally, signatures of endometriosis-associated endometrial-type epithelial clusters were enriched in ovarian cancers, reinforcing the epidemiologic associations between these two diseases.

## Introduction

Endometriosis is characterized by endometrial-like tissue growing outside of the uterine cavity, causing chronic pain, dysmenorrhea and infertility. Endometriosis is thought to occur in up to 10% of reproductive-aged people born with a uterus, globally impacting around 85 million people. True prevalence of the disease in the general population is likely grossly underestimated as many patients are asymptomatic and when symptoms do exist, they tend to be variable and nonspecific (e.g., pelvic pain). Consequently, a definitive diagnosis requires surgery with pathology confirmation (Hickey et al., 2014). In addition to pain and infertility, endometriosis is associated with increased risk of epithelial ovarian cancer, particularly tumor with clear cell and endometrioid differentiation (Lu et al., 2015; Pearce et al., 2012; Kohl Schwartz et al., 2017). Although clear cell ovarian cancer is relatively rare, these tumors are the second leading cause of ovarian cancer deaths due to their poor responses to standard chemotherapy regimens (Anglesio et al., 2011).

Many basic questions about endometriosis etiology remain unanswered. Endometriosis is likely a heterogenous disease entity, but efforts to subcategorize based on staging (which largely tracks overall disease burden), disease site, or infiltrative behavior have failed to identify meaningful correlates with pathologic features, severity or pathologic features (Zondervan et al., 2020). Endometriosis can be broadly categorized into ovarian endometriosis (endometrioma), superficial peritoneal endometriosis, deep infiltrating endometriosis (defined clinically as lesions that infiltrate > 5 mm under the peritoneal surface) and visceral endometriosis. Endometriomas exhibit a distinct cystic structure and almost exclusive occurrence within the ovary. Superficial lesions are patches of endometriosis on the lining of the peritoneum and can vary in color from clear, white and red to darker lesions that are brown, black or blue-ish in appearance. Deep infiltrating endometriosis is characterized by cancer-like local invasive behavior *and* frequent mutations in known cancer driver genes (Anglesio et al., 2017; Lac et al., 2019) but somewhat paradoxically, this subtype of endometriosis has comparatively modest associations with ovarian cancer risk; whereas ovarian endometrioma has been shown to enhance risk (Saavalainen et al., 2018).

Diagnostic criteria for endometriosis include the presence of endometrial-type epithelium and/or endometrial-type stroma in ectopic locations, often accompanied by hemosiderin-laden macrophages. Many lesions are microscopic, rendering traditional bulk genomic characterization approaches challenging as they are sensitive to isolation of uniform/pure specimens. In addition, endometriosis is often treated with ablation, which destroys the tissue. As such, the global molecular profiles of endometriosis lesions remain uncharacterized. Excision of endometriosis lesions, not only offers better outcomes for patients (Pundir et al., 2017) but enables histopathologic confirmation of suspected diagnoses and provides the opportunity to explore correlations of genotype and phenotype among diverse endometriosis lesions. To circumvent the challenges posed by the inter- and intra-patient cellular heterogeneity of endometriosis, we applied single cell RNA-sequencing (scRNA-seq) to 25 endometriosis lesions and endometriomas from seven patients and ovary specimens and eutopic endometrial samples from four women, including two patients not affected by endometriosis. We used these data to create a cellular atlas of endometriosis and to identify the molecular hallmarks of endometrial-type epithelial and stroma cells in the context of eutopic endometrium, endometrioma or extra-ovarian endometriosis. Finally, we leveraged the profiles of endometrial-type epithelium to deconvolute bulk expression profiles of clear cell and endometrioid ovarian cancer.

## Results

### Surgical and pathologic characterization of human endometriosis

To create a cellular atlas of endometriosis, we assembled a cohort of 9 endometriosis patients and 2 patients without endometriosis who underwent minimally invasive gynecologic surgery at our institution (Table 1). From these patients we collected a total of 37 specimens, which included 24 extra-ovarian (peritoneal) endometriosis specimens collected from 7 patients, 7 ovarian endometriomas from 6 patients, 4 eutopic endometrium samples (2 from endometriosis patients and 2 from patients without endometriosis) and 2 ovary tissues from women unaffected by endometriosis. Patients 7-9 and 11 had both endometriomas and endometriosis lesions profiled (Figure 1, Figure S1, Table 1). Both endometriosis-free patients were post-menopausal, both post-menopausal patients and one of the pre-menopausal patients were taking exogenous hormones at the time of surgery (Tables S1).

**Figure 1.**
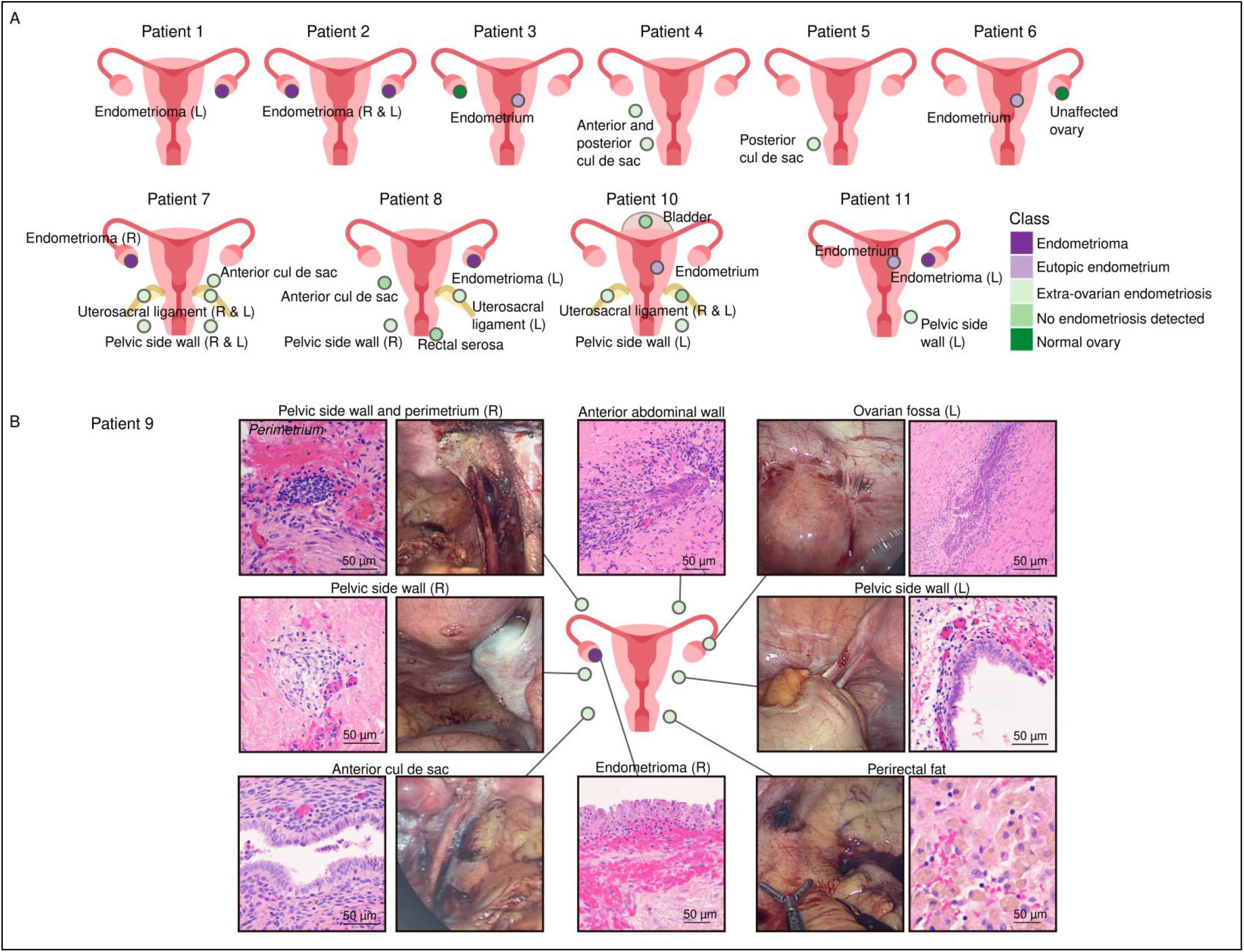
A cellular atlas of human endometriosis. (A) Patient cohort and specimens profiled. (B) Histologic and macroscopic features of specimens from patient 9.

A detailed pathology review was performed by a specialist gynecologic pathologist (Table 1, Figure 1A). A pathologic confirmation of endometriosis was defined as the presence of endometrial-type epithelium with endometrial-type stroma, the presence of endometrial-type stroma only or endometrial-type epithelium in association with hemosiderin pigmentation. All 7 the ovarian endometriomas had endometrial-type epithelium and stroma, plus hemosiderin present (Table S2). For the extra-ovarian endometriosis, 5 out of 24 specimens were suspicious for endometriosis during surgery, but had no endometriosis detected (NED) upon pathologic review (patient 8 - anterior cul de sac and rectal serosa; patient 9 - perirectal fat; patient 10 - bladder and left uterosacral ligament). To determine whether endometriosis was present in deeper levels, we examined 5 deeper sections cut at 50 μm intervals. Endometriosis was absent from all sections from patients 8 and 10, but endometrial-type stroma was detected in deeper levels for the perirectal fat specimen from patient 9. Of the remaining 16 specimens, 12 had epithelium, stroma and hemosiderin observed in at least one part. The remaining 4 specimens had two diagnostic components present in each. Endometrial-type stroma was present in all extra-ovarian lesions (Table S3).

### A single-cell atlas of endometriosis

To map the cellular and transcriptional features of endometrioma and endometriosis, we profiled these specimens using droplet based single cell RNA-sequencing (RNA-seq). In five instances, specimens with low cell yields were combined with specimens from similar anatomic locations in the same patient, to give a total of 32 samples sequenced - 7 endometriomas, 19 extra-endometriosis lesions, 4 eutopic endometrium specimens and 2 normal ovary tissues. The ovary specimen from patient 3 and left pelvic side wall specimen from patient 10 failed to meet quality control thresholds and were removed from the subsequent analyses (see Methods). A total of 257,255 individual cells were profiled, with a total of 6,606,535,712 reads sequenced (Table S4). After filtering out doublets, any cells with >20% mitochondrial transcripts or <200 genes detected, 207,949 cells remained for analysis. The median number of captured cells was 7,498 for eutopic endometrium (range = 3,408-20,833), 6,265 for endometrioma (range = 1,112-14,943), 5,038 for extra-ovarian endometriosis (range = 1,249-11,673). The observed cell number correlated with the targeted cell number (p=0.001, Spearman’s r=0.65), and was not significantly different between groups (p=0.084, one-way analysis of variance) (Figure S2A,B). Numbers of reads and genes per cell were also not significantly different between groups and did not differ based on whether specimens were cryopreserved prior to capture and sequencing (Figure S2A-D).

ScRNA-seq data were integrated using Harmony, regressing out the effects of inter-patient variability, sequencing batch and mitochondrial RNA content (Korsunsky et al., 2019) (Figure 2A-E). Cell cycle genes were not present in any of the first 20 principal components and so cell cycle regression was not performed. We identified 96 clusters and developed a systematic pipeline for cell-type assignment (see Methods and Figure S2E). First, we performed differential gene expression analysis to identify genes overexpressed in each cluster relative to all other clusters (log_2_ fold change = 0.2; p < 0.05) (Figure S2F and S2G). We then defined a set of rules for cell-type identification based on known hierarchies of canonical cell-type specific markers, for example *ACTA2* in the absence of other fibroblast markers denotes smooth muscle cells, but when co-expressed with fibroblast markers denotes activated fibroblasts (Methods, Figure S2E). For 84 out of 96 clusters we were able to assign cell identities using this approach (Figure 2B-E). For the remaining 12 clusters that did not overexpress canonical marker genes for any cell type, we calculated pairwise comparisons of all clusters and assigned cell identities based on the most correlated cluster (Figure S2H). Correlation values for cell type assignment ranged from 0.77-0.97 (Pearson’s correlations) and were significantly higher compared to random pairwise correlations (average random correlation r=0.019; Figure S2I), providing confidence that cell type assignment can be achieved using this approach.

**Figure 2.**
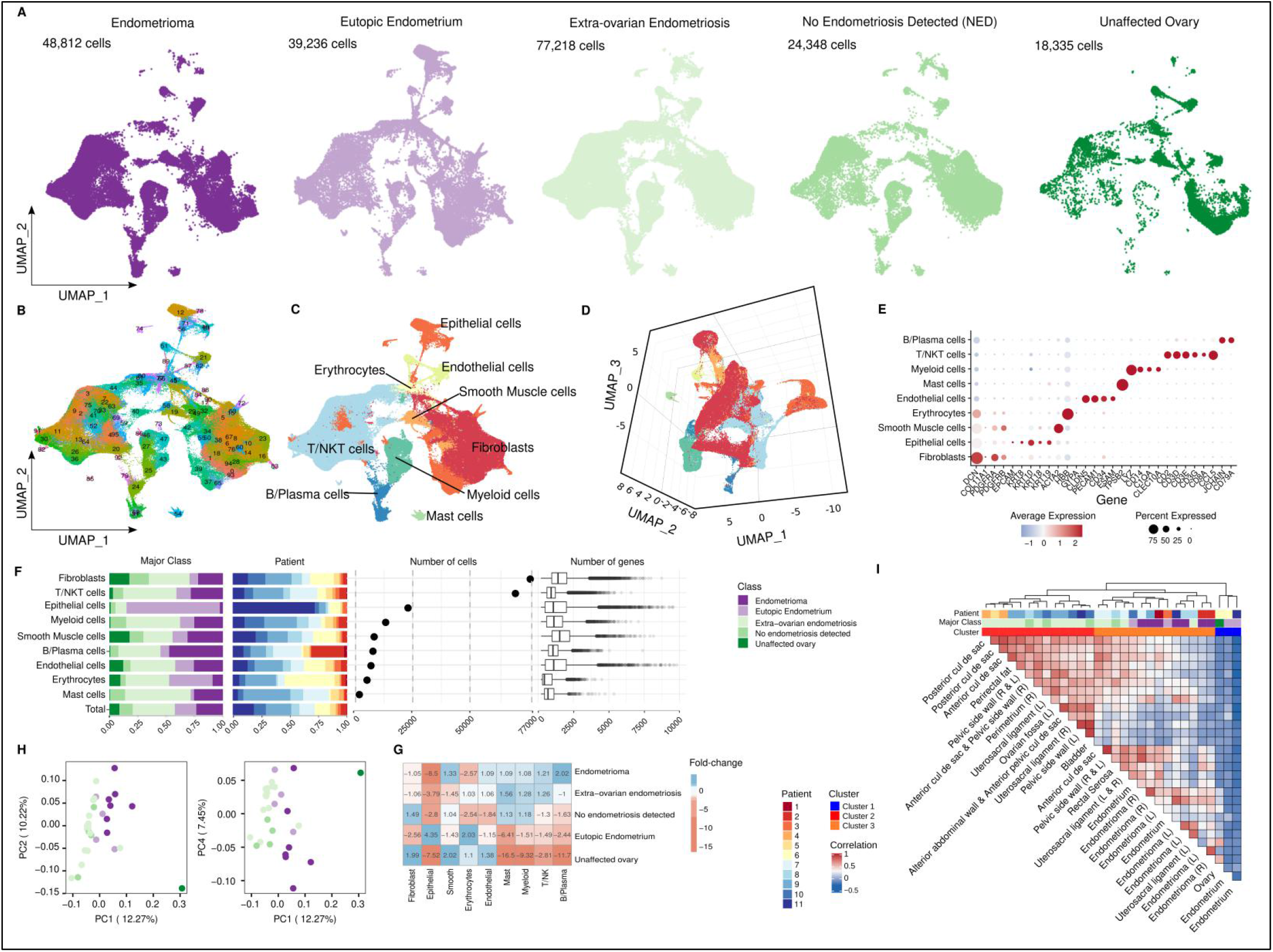
The cellular landscapes of endometrioma, peritoneal endometriosis, unaffected peritoneum, eutopic endometrium and unaffected ovary. (A) Uniform Manifold Approximation and Projection (UMAP) visualization of all sequenced cells (after filtering for quality) from 32 samples representing five major tissue-type classes. (B) UMAP plot with 96 clusters (harmony reduction using 25 dimensions and resolution of 3). (C) Major cell types identified, UMAP representation. (D) Three-dimensional UMAP representation. (E) Expression of representative markers across 9 major cell types. (F) Representation of each major class within each major cell type group, contribution of each patient to each group, frequencies of each cell type and number of genes detected in each cell type. ‘Total’ column represents the proportion of each patient or class in the major cell type overall, under the null hypothesis of no enrichment in a specific cluster. Box and whisker plots, boxes denote the interquartile range, bar denotes median number of genes detected per cell. The limits of the whiskers represent 1.5 * IQR (interquartile range) and outlier cells are indicated with individual dots. (G) Fold enrichment and depletion of each cell type across the five classes. (H) Principal component analysis. (I) Correlation, based on cell type frequencies, across all specimens profiled by scRNA-seq (Pearson correlation). We used the agglomeration method “complete” and “Euclidean” distance as clustering parameters.

Fibroblasts and stromal cells, identified by expression of *FAP*, *COL1A1* and *PDGFRA/B*, were the most abundant cell type present (n= 74,111 cells, 35.64% of cells remaining after filtering) (Figure 2E-F). T-cells/NKT-cells were the second most prevalent cell type present, comprising 69,703 (33.5% of cells). Keratin (*KRT7*, *KRT8*, *KRT10*, *KRT18*, *KRT19*) or *EPCAM*-positive epithelial cells (n=22,772 cells) represented 10.9% of the total population (Figure 2E-F). Other less common cell types included myeloid cells (n= 13,003 cells, 6.25% of the total population), smooth muscle cells (n=7,970 cells, 3.83% of the total population), endothelial cells (n=6,586 cells, 3.17% of the total population) and some erythrocytes that persisted after red blood cell depletion (n= 4,857 cells, 2.33% of the total population). Other immune cells included B and Plasma cells (n=7,452 cells, 3.58% of total population) and mast cells (n=1,495 cells, 0.72% of total population). Fibroblasts and smooth muscle cells had the greatest number of genes detected (average = 1,571.6 and 1,703.4 genes/cell), and as expected, erythrocytes had the lowest number of genes detected per cell (729.8 genes/cell). We compared the frequency of each cell type across the major tissue-type classes - endometrioma, extra-ovarian endometriosis, eutopic endometrium, tissue with no evidence of endometriosis and unaffected ovary, to identify deviations from the null distribution (the overall proportion of each cell type in the data set) (Figure 2G). Eutopic endometrium tissues were enriched 4.3-fold and 2-fold for epithelial cells and erythrocytes, respectively (p=2.2×10^−16^ and p=2.2×10^−16^, respectively, Chi-squared Test), and the large number of epithelial cells assigned to patient 11 (captured cell number = 38,400 after filtering) only partially explained the epithelial cell enrichment (after removing patient 11 eutopic epithelial cells the enrichment is 2.15 fold). Endometrioma tissues were depleted 8.5-fold for epithelial cells but enriched 2-fold for B and plasma cells. Extra-ovarian endometriosis was enriched 1.6-fold for mast cells, 1.3-fold for myeloid cells and 1.3-fold for T/NK-T cells (Figure 2G). Overall, each class of tissue type had a significantly different composition of cell types compared to the other four classes (p=2.2×10^−16^, Chi-squared Test).

We examined the overall relationships between specimens. In principal component analysis (PCA) endometriomas formed a group largely separated by PC1 (10.22% variance explained) (Figure 2H, Figure S2J) with separation of extra-ovarian endometriosis becoming evident in PC4 (7.45% of variance explained). We calculated pairwise correlations between all the specimens in the cohort based on the cell-type composition and performed unsupervised clustering. Samples separated into 3 major clusters (Figure 2I), Cluster 1 contained the normal ovary specimen and 2 of 4 eutopic endometrium samples. Cluster 2 contained the majority of the extra-ovarian endometriosis specimens (10 of 14) plus 3 of 4 specimens where no endometriosis was detected. All 7 endometrioma specimens were found in Cluster 3, along with 4 of 14 extra-ovarian endometriosis samples, 2 eutopic endometrium samples and the remaining specimen with no endometriosis detected (rectal serosa from patient 8). In some instances, samples from the same patient were highly correlated. For example, in patient 8, a left endometrioma was highly correlated with a nodule on the left uterosacral ligament (R^2^=0.77, Pearson correlation) and bilateral endometrioma samples from patient 2 were highly correlated (R^2^=0.83, Pearson correlation). In patient 9, all the extra-ovarian lesions were tightly correlated and were found within cluster 2, whereas the endometrioma from this patient was in cluster 3 and bore the most similarity to an endometrioma in patient 7 (R^2^=0.83, Pearson correlation).

### Epithelial components of eutopic and ectopic endometrium

Epithelial and stromal cells are the two major structural cell types present in endometriosis lesions, and so we characterized the heterogeneity within these populations. First, we isolated the 6,850 epithelial cells and re-clustered to identify 16 clusters that exhibited different patterns of canonical marker gene expression and were differentially represented across the tissue types (Figure 3A-C). Six of these were clusters of endometrial-type epithelial cells (EnEpi_1 to 6) that exhibited heterogenous expression of hormone receptors *ESR1* and *PGR* and six were mesothelial cell clusters (Meso_1 to Meso_6) expressing *WT1* and/or other mesothelial cell markers *PDPN, DES* and *CALB2*. Four clusters did not express canonical epithelial or mesothelial markers and were denoted as ‘unclassified’ epithelial clusters (UnEpi). UnEpi_1 was mostly derived from eutopic endometrium and expressed high levels of hemoglobin genes (*HBB*, *HBA1* and *HBA2*), low *PGR* and *ESR1* as well as differentiated epithelial cell markers (*TFF3* and *SLPI*; Table S5) and may represent luteal-phase endometrial-type epithelium. UnEpi_2 only had one defining marker (*C11orf96*; Table S5). UnEpi_3 and _4 were rare populations that represent possible immune cell contamination (Table S5).

**Figure 3.**
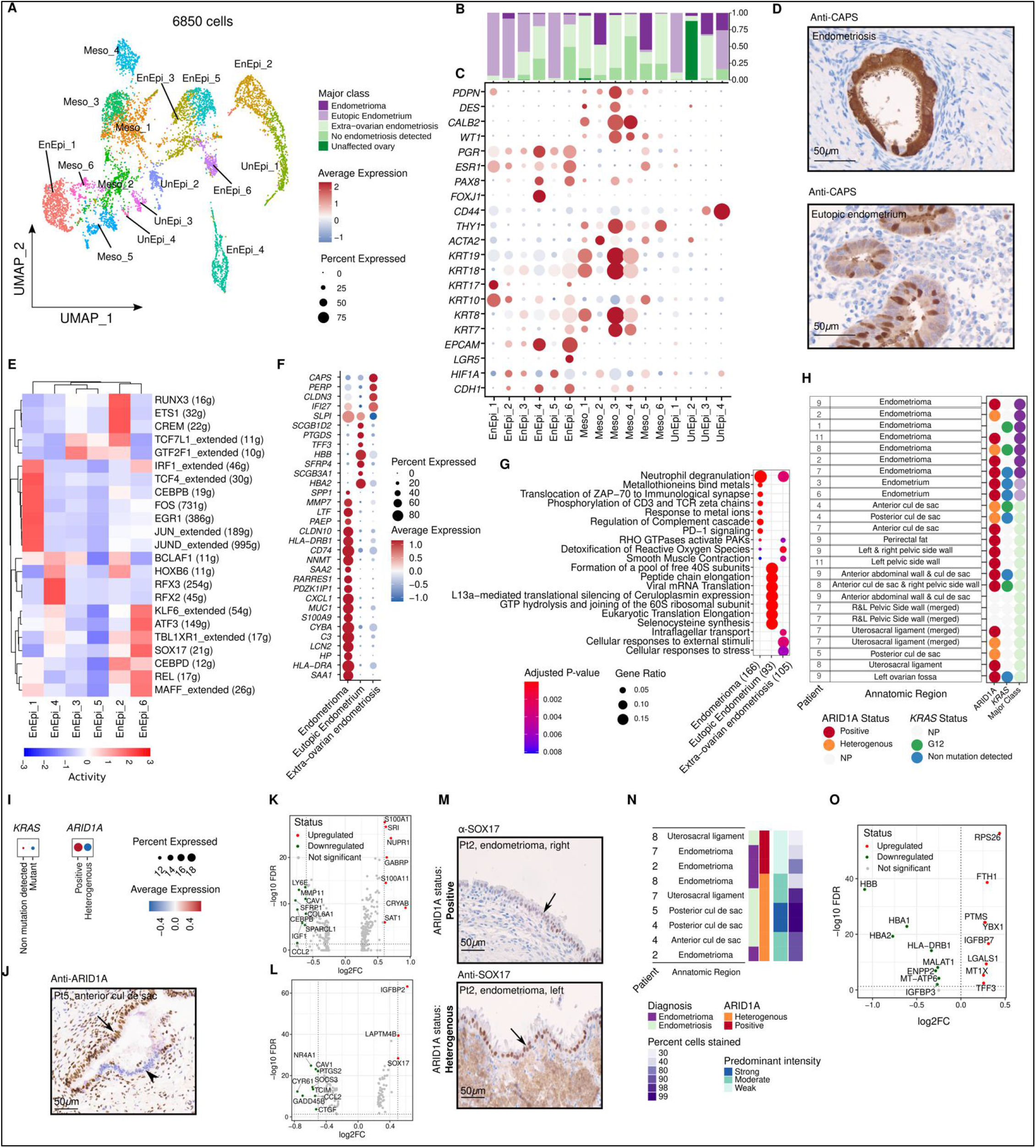
Epithelial components of eutopic endometrium and endometriosis. (A) UMAP of all epithelial cells, (B) frequency of each cluster by class. The total column represents the distribution of cells or class in all epithelial cells. (C) Marker gene expression and pathway analysis. (D) CAPS expression, immunohistochemical staining of eutopic endometrium and endometriosis. (E) Transcription factor regulon analyses using SCENIC. (F) Differential gene expression in endometrial-type epithelium in the context of endometrioma, eutopic endometrium or extra-ovarian endometriosis (pval < 0.05 and log_2_ FC > 1) (G) Pathway analysis, endometrial-type epithelium in the context of endometrioma, eutopic endometrium or extra-ovarian endometriosis. (H) Summary of ARID1A staining status and *KRAS* mutations detected in each lesion. NP, not profiled due to insufficient epithelial material available in the specimen. (I) Expression of ARID1A and *KRAS* mRNA by mutation state. (J) ARID1A immunostaining in a representative endometriosis lesion with heterogenous staining, positive staining for ARID1A is shown with the black arrow, negative epithelium is shown with the arrowhead. Posterior cul-de-sac lesion from Patient 5 is shown. (K) Differential gene expression in *KRAS* mutant versus wild-type endometrial-type epithelium (P < 0.05 and log_2_ FC = 0.6). (L) Differential gene expression in ARID1A heterogeneously staining versus positive endometrial-type epithelium (P < 0.05 and log_2_ FC = 0.5). (M) SOX17 staining in ARID1A positive and ARID1A heterogenous staining endometriomas from patient 2. (N) Summary of SOX17 staining by ARID1A staining status. (O) Differential gene expression in endothelial cells associated with positive or heterogenous ARID1A staining in endometrial-type epithelium.

The six clusters of endometrial-type epithelial cells could be broadly categorized based on keratin expression: EnEpi_1, 2, 4 and 6 expressed *KRT10* and/or *KRT17*, with EnEpi_6 also co-expressing low levels of *KRT8* and *KRT18*. EnEpi_3 and 5 exhibited low but pervasive expression of *KRT8* and *KRT18*. EnEpi_1, 2 and 5 were predominantly cells from endometriosis, with 93%, 88% and 97% of cells in these clusters derived from ectopic endometrium. The top pathways enriched in these clusters were distinct, with EnEpi_1 associated extracellular matrix remodeling, EnEpi_2 with immune cell interactions, and EnEpi_5 exhibiting a strong secretory cell signature. EnEpi_3, a second secretory cluster, were mainly derived from eutopic endometrium (56%) and 42% endometriosis (42%). EnEpi_4 and EnEpi_6 were *EPCAM* and *PAX8*-positive endometrial-type epithelial clusters that were detected in eutopic endometrium but were enriched 4.4 and 5.6-fold in peritoneal lesions. EnEpi_4 was a population of *FOXJ1+* ciliated epithelial cells enriched for the ‘cilium assembly’ pathway (P = 4.4×10^−20^) (Table S6). We validated EnEpi_4 by performing immunofluorescent staining of eutopic endometrium and endometriosis for CAPS, identifying CAPS-positive ciliated cells in both tissue types (Figure 3D). EnEpi_6 expressed high levels of *EPCAM*, *LGR5*, low expression of keratins plus intermediate levels of *PAX8*, *ESR1* and *PGR* and was enriched for cell-cell communication and splicing pathways (Figure 3A-C, Table S6). We note that endometrial-type epithelial cells were detected in all the four specimens that had no endometriosis upon pathologic review, ranging from 24 and 26 cells in the two NED specimens from patient 8, to 48 and 232 cells in the NED specimens from patient 10, suggesting that, particularly for the latter patient, the portion used for scRNA-seq may have contained endometriosis epithelium. We implemented SCENIC to identify the main transcription factors driving the transcriptional programs of each cluster (Figure 3E). Unsupervised clustering was performed based on similarity of transcriptional regulon activity. The first group contained EnEpi_1 which was characterized by activation of IRF1, TCF4, CEBPB and FOS regulons. EnEpi_4 was characterized by active RFX2, RFX3, HOXB6 and BCLAF1 regulons. EnEpi_3 and EnEpi_5 were characterized by GTF2F1 activity. EnEpi_2 also had activation of the GTF2F1 regulon as well as RUNX3, ETS1 and CREM. Finally, EnEpi_6 exhibited strong activation of KLF6, ATF3, SOX17, REL and MAFF.

Endometrial-type epithelium exhibited marked differences in gene expression in the context of endometrioma, eutopic endometrium or extra-ovarian endometriosis. The 220 genes overexpressed in endometrioma (log_2_ FC > 0 and adjusted p < 0.05) included progestagen associated endometrial protein (*PAEP*, FC=1.44, adjusted P = 9.9×10^−33^) and were enriched in pathways associated with immune cells interactions, including PD-1 signaling (P = 2.9×10^−6^), and hypoxic insult, as reflected by an enrichment of ‘Detoxification of Reactive Oxygen Species’ (P = 6.1×10^−3^) (Figure 3F,G, Table S7, Table S8). The latter pathway was enriched in both endometriomas and peritoneal endometriosis, as was RHO GTPase effectors (P = 9.2×10^−4^), neutrophil degranulation (P = 1.7×10^−10^) and cellular responses to external stimuli (P = 3.1×10^−3^), highlighting interactions with microenvironmental cells and stimuli as unifying hallmarks of endometriosis independent of location. Ciliated cell pathways and markers (including *CAPS*) were specifically enriched in peritoneal endometriosis, mirroring the enrichment of the EnEpi_4 ciliated epithelial cell cluster in this group. Finally, translation pathways reflecting the differentiated secretory epithelial clusters were enriched in eutopic endometrium.

Somatic mutations in “cancer driver” genes including *ARID1A* and *KRAS* (Anglesio et al., 2017; Lac et al., 2019; Suda et al., 2020) are known to occur in endometriosis, and so we sought to determine the transcriptional consequences of these mutations when they occur *in vivo*. Formalin-fixed, paraffin embedded tissue sections adjacent to the specimens used for scRNA-seq were available for 25 out of 32 specimens, of these 21 had sufficient epithelial content to quantify ARID1A expression using immunohistochemistry as a mutation surrogate and 10 specimens had sufficient epithelial content to successfully identify *KRAS* mutations by digital droplet PCR (ddPCR). A summary of the mutation profiling is shown in Figure 3H. The two endometrium specimens profiled were wild-type for both genes. Six patients had evidence of at least one somatic mutation in one or more endometriosis lesions. Two endometriomas and four extra-ovarian lesions exhibited heterogenous ARID1A expression indicative of heterozygous *ARID1A* loss of function mutations. Two of the five endometriomas and two of the five extra-ovarian lesions subjected to *KRAS* genotyping harbored mutations at codon 12. *ARID1A* and *KRAS* expression were highest in EnEpi_4 and _6, the two clusters enriched in endometriosis specimens (Figure S3C). When we compared expression by mutation status, *ARID1A* mRNA expression was 2.1-fold lower in specimens where endometrial-type epithelium exhibited heterogenous protein staining compared to cases with strong ARID1A staining (Figure 3I,J; Figure S3E,F). By contrast, *KRAS* gene expression was only marginally elevated (by around 5%) in mutant compared to wild-type cells (Figure 3K). We then performed differential expression analysis to identify genes (Figure 3K, L) and pathways (Figure S3G,H) associated with mutation of *KRAS* or *ARID1A*, after removing genes associated with class to minimize the impact of this confounding variable. *KRAS* mutation was associated with 302 differentially expressed genes (log_2_ FC > 0, adjusted P < 0.05) including known *KRAS* target gene *S100 calcium-binding protein A1* (*S100A1*; log_2_ FC= 0.61, adjusted P = 9.16×10^−33^) and transcriptional regulator *nuclear protein 1* (*NUPR1*; log_2_ FC= 0.70, adjusted P = 2.05×10^−29^)(Figure 3K, Table S10). 118 genes were differentially expressed in endometrial-type epithelial cells associated with heterogenous ARID1A staining compared to those with homogenous positive staining (log_2_ FC > 0, adjusted P < 0.05)(Figure 3L, Table S11). The most upregulated gene associated with *ARID1A* loss of function was known ARID1A target gene *IGFBP2* (log_2_ FC= 0.62, adjusted P = 2.24×10^−68^) (Suryo Rahmanto et al., 2020). Additional targets included *lysosome-associated protein transmembrane-4β* (*LAPTM4B*; log_2_ FC= 0.51, P = 1.24×10^−44^) and *SRY-box 17* (*SOX17*; log_2_ FC= 0.50, adjusted P = 1.51×10^−33^, Figure 3L). We have recently identified SOX17 as a novel marker of secretory fallopian tube epithelia and high-grade serous ovarian cancer, where it positively regulates angiogenesis (Chaves-Moreira et al., 2020; Dinh et al., 2021; Reddy et al., 2019). SOX17 protein expression was validated in the same tissues using immunohistochemistry performed on 6 specimens with heterogenous ARID1A staining and 2 specimens with positive ARID1A staining. The proportion of endometrial-type epithelial cells expressing SOX17 was higher in lesions with heterogeneous ARID1A staining. Heterogeneous ARID1A staining was also associated with moderate/strong SOX17 staining, whereas lesions with positive ARID1A staining exhibited weak SOX17 staining (Figure 3M & N). We therefore interrogated the endothelial cell compartment associated with ARID1A mutation status and found that endothelial cells proximal to ARID1A-mutant endometriosis lesions up-regulate expression of a select handful of genes including *ferritin heavy chain* (*FTH1*, log_2_ FC= 0.29, adjusted P = 8.18×10^−44^), *parathymosin* (*PTMS*, log_2_ FC= 0.27, adjusted P = 1.79×10^−29^), *Y box binding protein 1* (*YBX1*, log_2_ FC= 0.27, adjusted P = 2.04×10^−29^), *galectin-1* (*LGASL1*, log_2_ FC= 0.28, adjusted P = 1.58×10^−14^) and *metallothionein 1X* (*MT1X*, log_2_ FC= 0.25, adjusted P = 1.85×10^−10^)(Figure 3O).

### Endometriomata impact mesothelial differentiation

Five epithelial clusters were defined as mesothelial due to expression of canonical mesothelial cell markers (*WT1*, *CALB2*, *DES* and/or *PDPN*). Mesothelia are a specialized type of epithelium that lines the peritoneum, the pleurae and pericardium (Figure 3B,C). Modified peritoneal mesothelium also covers the ovary as a monolayer termed the ovarian surface epithelium, which can also become trapped within ovarian inclusion cysts that develop following ovulation (Auersperg et al., 2001). We examined the pseudotime relationships between mesothelial cells clusters with Monocle3 (Qiu et al., 2017). Ovarian mesothelial cells associated with endometrioma were enriched at an earlier timepoint in the trajectory and exhibited lower expression of differentiated markers compared to mesothelial cells from unaffected ovary tissue (Figure 4A). Peritoneal mesothelium sat later in the pseudotime space and was more differentiated than unaffected ovary or endometrioma-associated mesothelium. Meso_5 was derived from endometrioma and peritoneal lesions (both with and without pathologic confirmation of endometriosis). This cluster expressed *PDPN* and *WT1* but not *CALB2* or *DES*, and was the only mesothelial cluster that expressed *ESR1* and *PGR*. Meso_5 expressed *ACTA2* and also exhibited a markedly different keratin profile, expressing *KRT10* but not *KRT7*, *KRT8*, *KRT18* and *KRT19*, the dominant keratins in the differentiated mesothelial clusters (Figure 3B,C). This suggests that mesothelial cells associated with endometrioma tend to exhibit a modified uncommitted mesothelial-mesenchymal phenotype. Novel markers overexpressed by endometrioma-associated mesothelial cells included *IGFBP1, IGFBP2, IGFBP3, RBP1*, *MEG3* and *SOX4* (Figure 4B, C). Endometrioma-associated mesothelial cells exhibited lower expression of markers including *MSLN*, *SLPI*, *PRG4*, *ADIRF* and *TFPI2* when compared with mesothelial cells associated with extra-ovarian endometriosis (Figure 4A-C).

**Figure 4.**
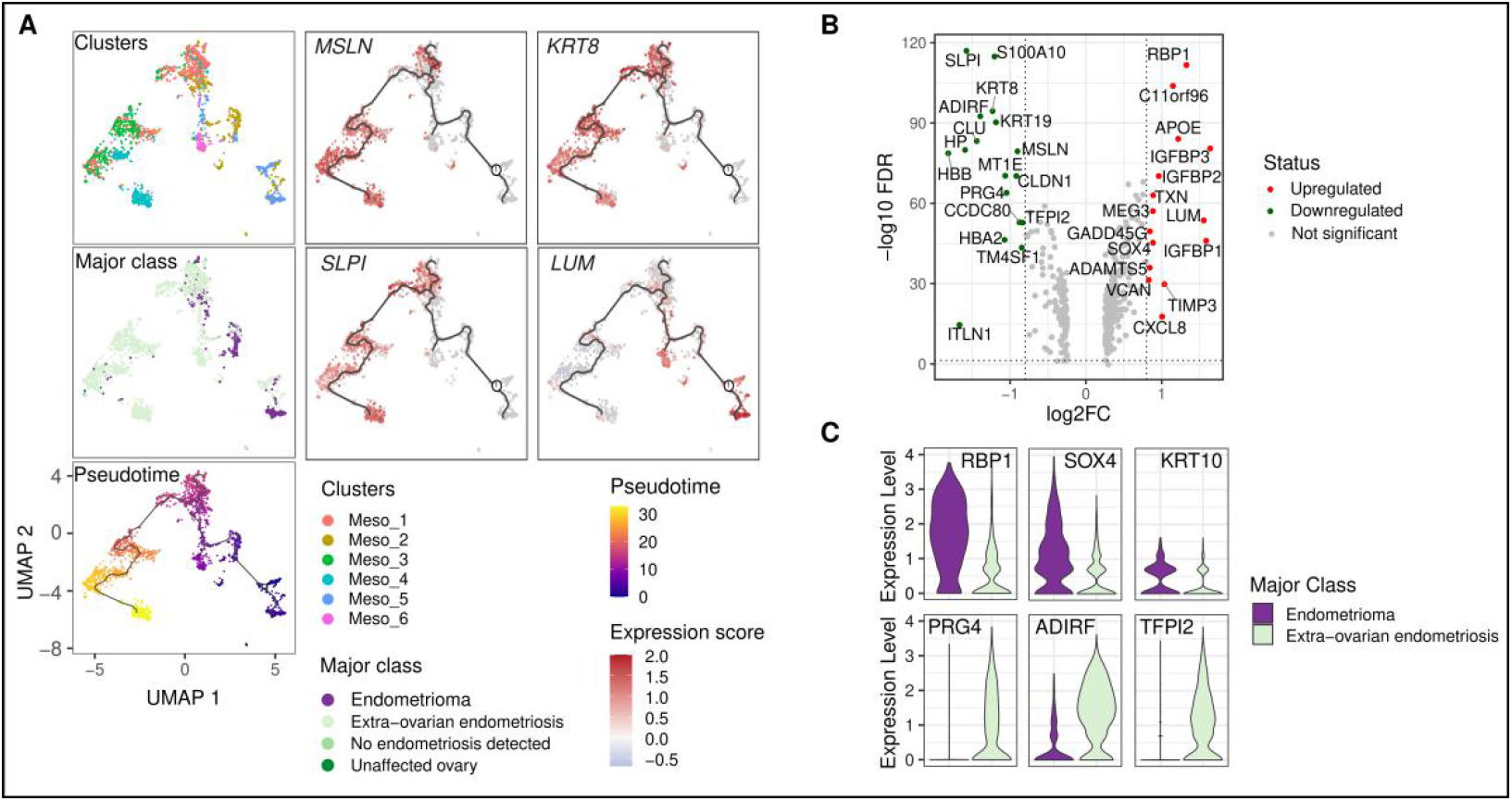
Endometriosis affects mesothelial differentiation. (A) Pseudotime analysis for mesothelial cells (B) Differential gene expression in mesothelial cells adjacent to endometrioma compared to extra-ovarian endometriosis (labelled genes are those where adjusted P < 0.05 and absolute log_2_ FC > 0.8). (C) *RBP1*, *SOX4* and *KRT10* are overexpressed in mesothelial cells proximal to endometriomata and *PRG4*, *ADIRF* and *TFPI2* overexpressed in mesothelial cells associated with extra-ovarian endometriosis.

### Mesenchymal components of eutopic endometrium, endometrioma and endometriosis

We turned our attention to the large population of mesenchymal cells in the data set. Sub-clustering of the 74,111 mesenchymal cells identified 20 distinct clusters that could be stratified into three major groups - ‘host’ organ ovarian fibroblasts (OvF), ‘host’ organ peritoneal fibroblasts (PerF) and endometrial-type stroma (EnS) (Figure 5A-D, Table S11). Clusters OvF_1 and OvF_2 were largely exclusive to the unaffected ovary (Figure 5A). Cluster OvF_1 was the largest fibroblast cluster (12,910 cells), exhibiting relatively low levels of canonical fibroblast markers *DCN* and *COL1A1*. Genes over/under-expressed in this cluster were associated with protein production and translation (Table S12). OvF_2 was a rare cluster (186 cells) with high expression of *DES* and *ACTA2* (Figure 5B,E). Peritoneal fibroblasts comprised 11 clusters that tended to have higher expression of *THY1*, *COL1A1* and *DCN* compared to ovarian and endometrial fibroblasts (Figure 5E). Peritoneal fibroblasts could be broadly categorized into *PDGFRA+* clusters (PerF_3, 4, 5, 9 and 10), *PDGFRB+* clusters (PerF_1 and 2), dual *PDGFRA+/PDGFRB+* clusters (PerF_7), and *PDGFRA/B* weak/absent (PerF_6, 8 and 11).

**Figure 5.**
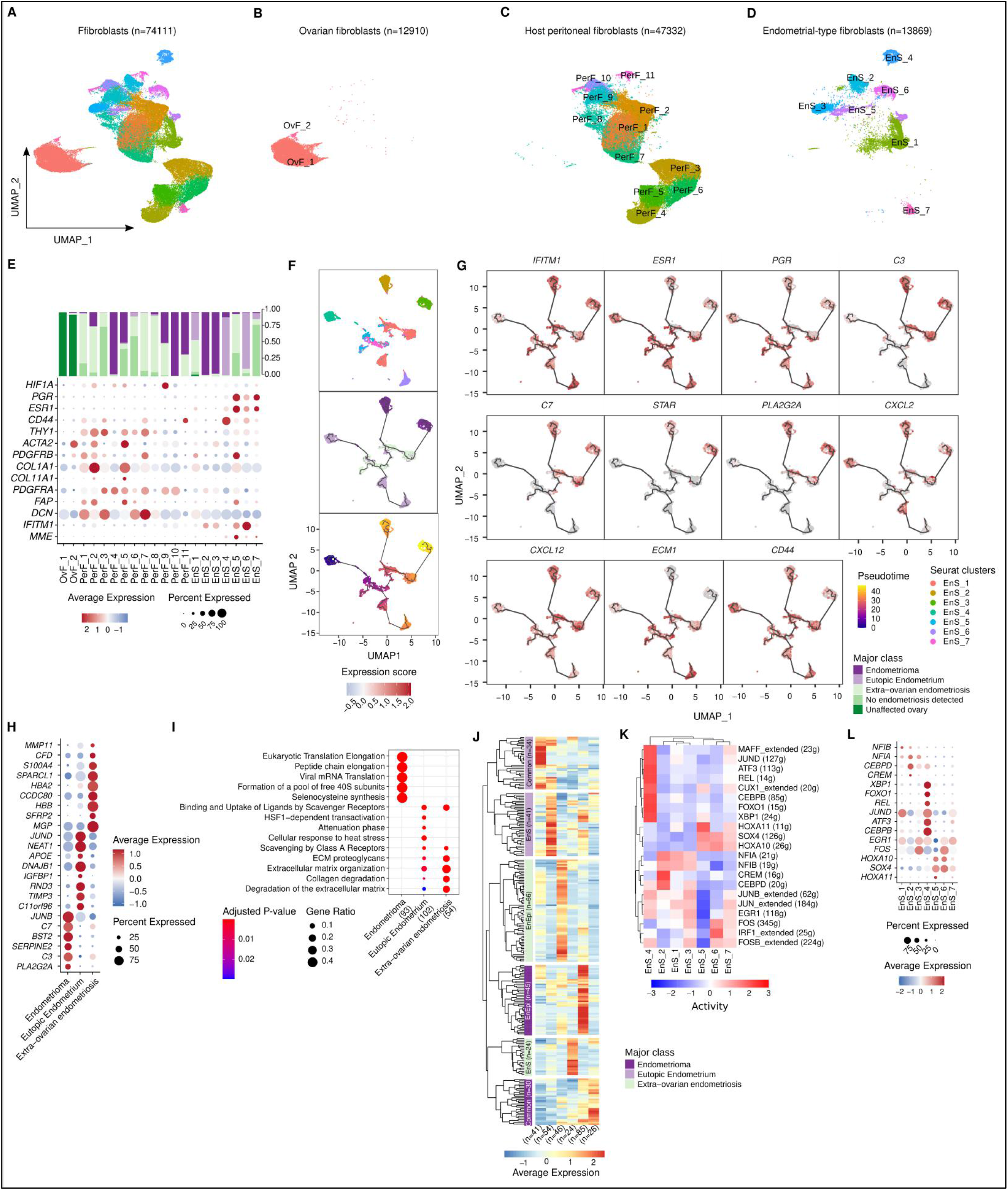
Signatures of endometrial-type stroma and mesenchymal cells associated with endometriosis. UMAP of mesenchymal cells in (A) all tissue types (B) unaffected ovary from an endometriosis-free patient, (C) peritoneal fibroblasts (D) endometrial-type stroma, (E) Marker gene expression and cluster frequencies. (F) Pseudotime analysis performed using Monocle3, and (G) marker gene expression. (H) Differential gene expression in endometrial-type stroma in the context of endometrioma, eutopic endometrium or extra-ovarian endometriosis (log_2_ fold change ≥ 0.8, P < 0.05) (I) Pathway analysis, endometrial-type stroma in the context of endometrioma, eutopic endometrium or extra-ovarian endometriosis. (J) Heatmap, coordinate expression of genes in EnEpi and EnS across eutopic endometrium, endometrioma and extra-ovarian endometriosis. Ordering of rows and columns (genes) are supervised, genes ranked in order of decreasing expression within each class. (K) Transcription factor regulon analyses using SCENIC. (L) Differential expression of candidate TFs across EnS clusters.

Seven clusters were identified as endometrial-type stroma (EnS) due to expression of known markers of endometrial stromal cells *MME* and or *IFITM1*, and/or predominant occurrence in eutopic endometrium (Figure 5E). We implemented Monocle3 to infer a pseudotime trajectory rooted in EnS_4, which expressed the highest levels of stem cell marker CD44 and was derived predominantly from eutopic endometrium (90%). This created a bifurcated trajectory (Figure 5F). Cluster EnS_6 represented one endpoint and was composed of differentiated endometrial-type stroma which exhibited the highest expression of *IFITM1* of all the EnS clusters, and moderate expression of *ESR1* and *PGR*. EnS clusters 1, 5 and 7 resided at intermediate pseudotime points in the trajectory. EnS_1 was the most abundant endometrial-type stroma cluster; 43% of cells in this cluster were derived from eutopic endometrium and 55% were derived from endometriosis. EnS_1 expressed *MME, PGR, ESR1* and *PDGFRB* and was enriched for heme-response pathways (Table S12). EnS clusters 5 (associated with extracellular matrix remodeling) and 7 (associated with stress response pathways) were predominantly composed of cells from peritoneal tissues (96 and 89%, respectively) and expressed the highest levels of *PGR* and *ESR1*. These two clusters were both observed in peritoneal lesions both with and without pathologic confirmation of endometriosis, again suggesting the portion subjected to scRNA-seq did contain endometriosis. Biomarkers expressed by EnS_5 include *progesterone receptor membrane component 1* (*PGRMC1*) and *receptor activity modifying protein 1* (*RAMP1*). EnS_5 and 7 both overexpressed matrix metalloproteinase-11 (*MMP11*). EnS clusters 2 and 3 represented alternative trajectory endpoints, both expressed *IFITM1* and modest expression of *PDGFRA* and *ACTA2* respectively (Figure 5F). Cells in these clusters (98 and 96% respectively) were derived almost exclusively from endometrioma samples. EnS_2 expressed an immunomodulatory set of genes including *CXCL12* and *CXCL2* (Figure 5G). Given the indication that endometrioma-associated EnS clusters exhibit more activation of pro-inflammatory pathways, we tested for gene signatures that differed across endometrial-type stroma by site. Endometrial-type stroma within endometriomas overexpressed *Phospholipase A2* (*PLA2G2A*), which when secreted plays a role in inflammation and neurological disorders. Innate immunity components complement proteins *C3* and *C7* were also overexpressed along with transcription factor *JUNB* and *bone marrow stromal antigen 2* (*BST2*) (Figure 5H, Table S13). The noncoding RNA *NEAT1*, transcription factor *JUND* and heat shock protein *DNAJB1* were abundantly expressed in endometrial stroma in eutopic endometrium but downregulated in endometriosis. Peritoneal endometriosis was associated with *matrix gla protein* (*MGP*), hemoglobin genes (*HBB* and *HBA2*) and tumor suppression and adhesion modulator *SPARC-like protein 1* (*SPARCL1*) (Figure 5H). At the pathway level, eutopic endometrium and extra-ovarian endometriosis were similar, with enrichment of extracellular matrix organization pathways (Figure 5I, Table S14). Endometrial type stroma within endometriomas was enriched for translation pathways (p=2.3×10^−65^ and 3.6×10^−64^), indicative of a secretory phenotype (Figure 5I). We noted overlap in the genes differentially regulated in endometrial-type epithelium and stroma associated with eutopic endometrium or endometrioma but not extra-ovarian endometriosis (Figures 3 and 5), for example, in endometriomas, *C3* and *C7* are highly expressed by both types and complement pathways were enriched (Tables S7, S8, S13 & S14) suggesting a coordinated transcriptional response occurs (Figure 5J).

Transcription factor regulon analyses identified an enrichment of FOXO1, XBP1, MAFF and JUND regulons in the putative endometrial-type stroma progenitors. Inflammatory EnS clusters (EnS_2 and EnS_3) and EnS_1 were associated with activation of pro-differentiation factors NFIA and NFIB (Chen et al., 2017) with EnS_2 showing high activation of the CREM regulon. EnS_3 shared activation of FOS, IRF1 and FOSB with differentiated endometrial stroma clusters EnS_6 and EnS_7. EnS_5-7 exhibited the highest expression of *PGR* and *ESR1* and were associated with HOXA10/11 and SOX4 activation (Figure 5K). *SOX4* expression was particularly high in EnS_5 and EnS_6 (Figure 5L).

### Deep and superficial peritoneal endometriosis involves transcriptional reprogramming in endometrial-type epithelium and mesenchymal cells

Surgical images were available for 24 out of 25 lesions and were reviewed by three expert minimally invasive gynecology surgeons, and consensus classifications of endometriosis subtype were assigned for extra-ovarian lesions (Table 1, Figure 1, Figure S1). Five lesions were categorized as deep infiltrating endometriosis and 8 were classified as superficial endometriosis (n=8). We asked whether single cell transcriptome profiles indicate that deep and superficial endometriosis are two biologically distinct entities. We integrated data sets for all 13 extra-ovarian samples to create a data set of 69,131 individual cells. We then simulated a background data set based on the frequencies of the different cell types in the actual data and measured Euclidian distance between deep and superficial lesions, based on frequencies of all cell types (Figure S4C). There was a modest suggestion that deep and superficial endometriosis were significantly different when we considered all cell types present (p=0.046, compared to a background of 1,000 randomly generated data sets, *pnorm* R function using mean and standard deviation from background) (Figure 6A). By contrast, hemorrhage and fibrosis were also not associated with cellular composition (p=0.56 and p=0.82, respectively).

**Figure 6.**
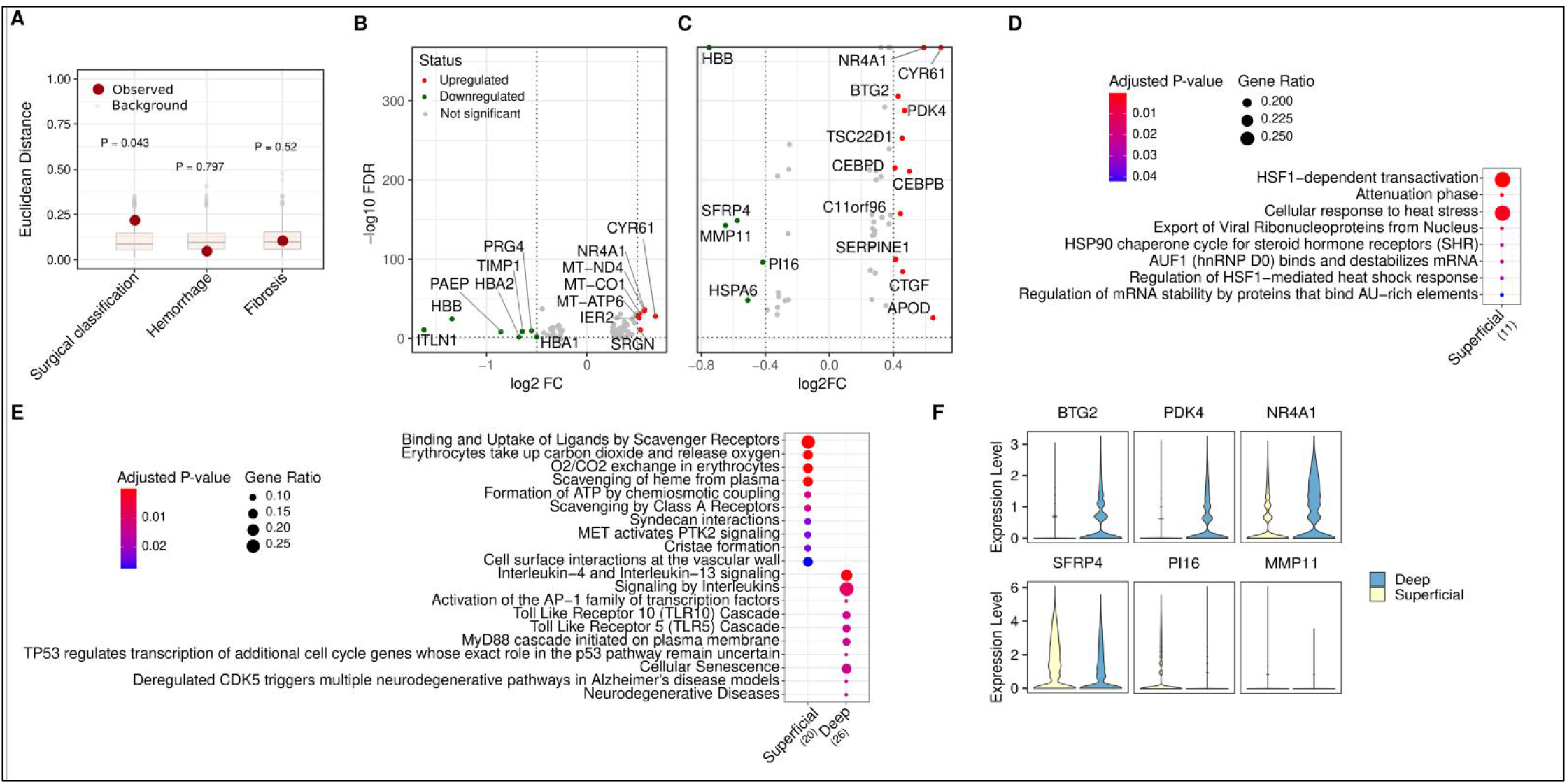
Molecular correlates of peritoneal endometriosis subtypes. (A) Euclidian distance between samples, compared to a simulated background distribution. Deep/superficial status, presence/absence of hemorrhage or fibrosis were all surveyed. (B) Differentially expressed genes (log_2_ fold change ≥ 0.5, P < 0.05) and (C) pathway enrichment in endometrial-type epithelial cells associated with deep or superficial endometriosis. (D) Differentially expressed genes (log_2_ fold change ≥ 0.4, P < 0.05) and (E) pathway enrichment in fibroblasts associated with deep or superficial endometriosis (F) Expression of *BTG2*, *PDK4* and *NR4A1* markers were enriched in deep endometriosis and expression of *SFRP4*, *PI16* and *MMP11* enriched in superficial endometriosis.

We posited that differences in deep and superficial endometriosis may more apparent in the expression signature of cells rather than overall cellular composition of lesions and so we tested for associations between endometrial-type epithelial or fibroblast gene expression and surgical annotations. Epithelial cells in deep infiltrating endometriosis were associated with upregulated expression of mitochondrial genes (*MT-ND4*, *MT-CO1* and *MT-ATP6*) and Nuclear Receptor Subfamily 4 Group A Member 1 (*NR4A1*) (Figure 6B,C; Table S15). *Intelectin-1* (*ITLN1*) was overexpressed in superficial lesions and both at the gene and pathway level, pathways associated with exposure to heme were enriched in superficial lesions (Figure 6B,C). Senescence and inflammatory pathways including interleukin-4 and −13 signaling were enriched in deep endometriosis (p=1.1×0^−2^ and p=4.3×10^−4^, Figure 6C). Differential gene expression in deep *versus* superficial lesions were more marked in fibroblasts compared to epithelial cells (although this is likely due in part to the larger number of cells). *NR4A1* was overexpressed in fibroblasts associated with deep endometriosis (log_2_ FC=0.58, p=4.1×10^−42^), plus *BTG2, CEBPB* and *CEPBD*, *SERPINE1* and YAP target gene *CTGF*. Fibroblasts associated with superficial lesions overexpressed *MMP11*, *SFRP4* and *PI16* (Figure 6D, Table S16). Stress associated transcription was upregulated in fibroblasts in superficial endometriosis, while no pathways were enriched in fibroblasts in deep endometriosis lesions (Figure 6E).

### Linking endometriosis signatures to endometriosis-associated ovarian cancers

Endometrioid and clear cell ovarian cancers are associated with a personal history of endometriosis, suggesting that endometrial-type epithelial cells may be precursors for these tumors. We implemented MuSiC (Wang et al., 2019) to test whether cluster-specific signatures of endometrial-type epithelium were enriched in these tumor types. Across three independent data sets, clear cell and endometrioid ovarian cancers (Tan et al., 2019; Tothill et al., 2008; Winterhoff et al., 2016) consistently showed a strong enrichment of signatures for ciliated cluster EnEpi_4, and to a lesser extent, cell-cell communication-associated cluster EpEpi_6 (Figure 7A-C). Both of these clusters of endometrial-type epithelial cells were enriched in endometriosis lesions relative to eutopic endometrium (Figure 3B). There was no evidence of enrichment for the signatures of endometrial-type epithelial cell clusters that were derived from eutopic endometrium.

**Figure 7.**
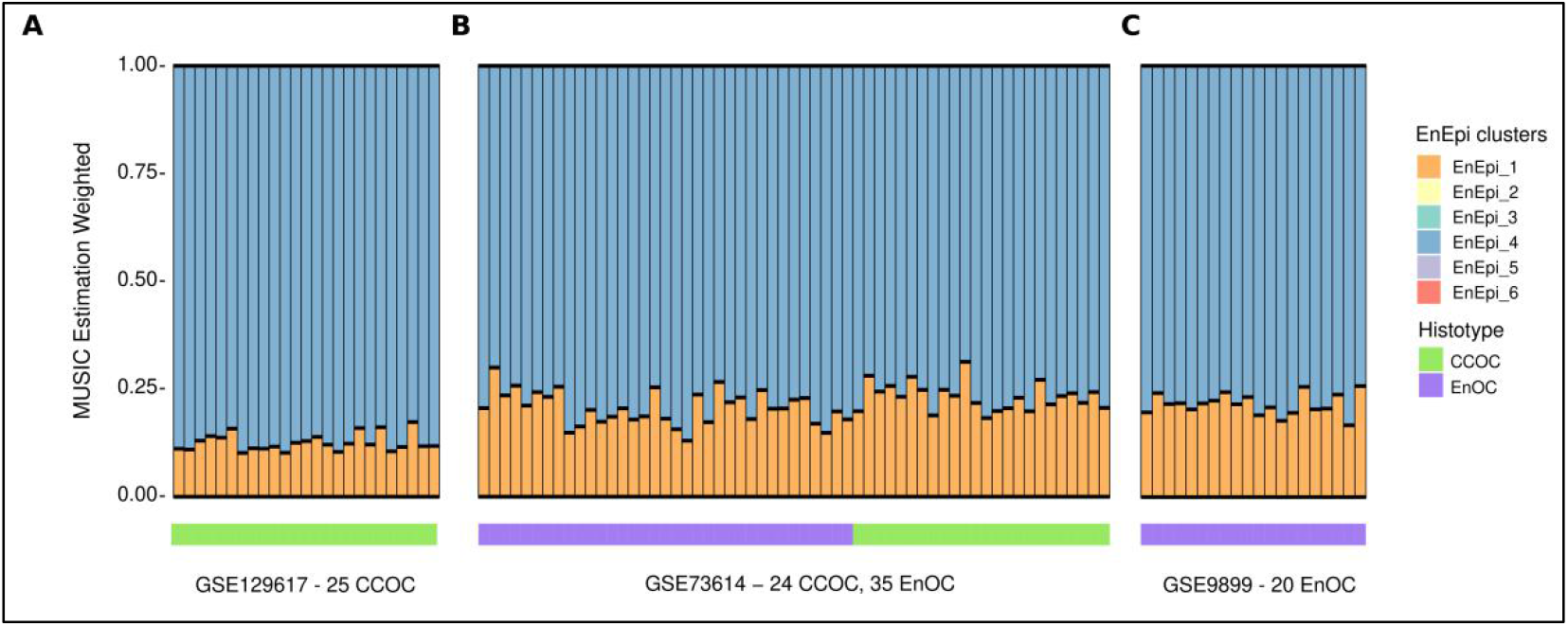
Deconvoluting endometriosis-associated ovarian cancers with single cell endometriosis signatures. Deconvolution of (A) 25 clear cell ovarian cancers (CCOC, 24 primary tumor specimens, one ascites sample) (B) 14 endometrioid ovarian cancers (EnOC) and (C) 24 CCOC and 35 EnOC tumors, based on signatures of 6 endometrial-type epithelial subclusters.

## Discussion

Endometriosis is a common but poorly studied condition of unknown etiology and poorly characterized pathogenesis. We generated a cellular atlas of endometriosis incorporating over 30 tissues from 11 patients, analyzing around 250,000 individual cells. In doing so we catalogued the epithelial component but also the stromal cells and immune cells in the microenvironment which are not passive bystanders but play active roles in endometriosis pathogenesis. We leveraged the data to ask some key clinical questions. First, these analyses add to a growing body of literature to suggest that endometriomas and peritoneal lesions are two distinct disease entities. Although heterogeneity across patients was observed, for some patients, cellular composition of lesions was highly correlated across sites. This composition may be influenced by underlying clonal expansion of endometriosis epithelium as it transits from endometrium and colonized ectopic sites where clonality has also been observed across lesions types through DNA-based analyses (Moore et al., 2020; Praetorius et al., 2021; Suda et al., 2018).

Despite the small size of most endometriosis lesions, in all except one endometriosis specimen we were able to capture the endometrial-type epithelial and stromal components that are diagnostic for the disease. First, we asked whether the molecular profiles of endometrial-type epithelium and stroma differ by site. In addition to observing striking differences in gene expression by context, we noted convergent expression of genes and pathways in endometrial-type epithelium and stroma in the context of endometrioma or extra-ovarian endometriosis and also in comparisons of deep and superficial peritoneal lesions, consistent with a recent multi-organ analysis of structural cells that highlighted pervasive organ-specific patterns of gene expression (Krausgruber et al., 2020). Both endometrial-type epithelium and endometrial-type stroma exhibited greater activation of immune pathways in the context of endometriomata, with complement proteins *C3* and *C7* expressed by both cell types, indicative of dysregulated innate immunity. This may be due to elevated apoptosis in the hypoxic microenvironment in an endometrioma, since cells undergoing apoptosis can activate the complement pathway, where dying cells opsonized by complement components help phagocytic cells such as macrophages dispose of the apoptotic cells. Moreover, serum and peritoneal fluid levels of C3 are elevated in women with endometriosis compared to controls (Hasan et al., 2019; Kabut et al., 2007), although endometrioma-specific analyses have not yet been performed. Interestingly, lower levels of the inactive iC3b are observed in both the serum and peritoneal fluid of endometriosis patients, yet higher levels of C3c and SC5b-9, possibly due to cleavage of iC3b by regulatory protein Factor I or complement receptor 3, releasing C3c leaving behind C3dg bound to cells. Bound C3dg can still interact with CR3/CD11b found on phagocytes to stimulate phagocytosis, and with CR2/CD21 found on B cells which can augment signaling through the B cell receptor (Ricklin et al., 2016). Whether the interaction of complement and CD21 on B cells contributes to the increase in autoantibodies seen in endometriosis patients is yet to be determined, but high expression of *BST2/CD317* and *CXCL12* on endometrial-type stroma associated with endometriomata may contribute to altered innate immunity in this context specifically.

It has been proposed that two subtypes of peritoneal disease exist - superficial peritoneal disease and deep infiltrating endometriosis (Brosens et al., 1993), and these categories are widely used in endometriosis research. We asked whether these subtypes were supported by the cellular and molecular profiles of peritoneal endometriosis categorized as deeply infiltrating or superficial. The overall cellular landscape of endometriosis did associate with deep/superficial status, but not fibrosis or hemorrhage, although we note we were powered only to detect very strong effects. We also observed that both endometrial-type epithelial cells and fibroblasts exhibited marked differential gene expression associated with deep or superficial status. These results may suggest that deep and superficial disease are not distinct entities but parts of the same disease continuum resulting from transcriptional reprogramming, consistent with recent genomic analyses (Praetorius et al., 2021). *NR4A1* was identified as an epithelial and fibroblastic marker of deep endometriosis and has been previously implicated in endometriosis-associated fibrosis (Zeng et al., 2018). *NR4A1* has been proposed as a therapeutic target for endometriosis (Mohankumar et al., 2020), although the precise functions of their protein in disease pathogenesis have yet to be elucidated. NR4A orphan nuclear receptors facilitate transcriptional and posttranscriptional responses to changes in the cellular microenvironment and may therefore serve as a hub for the altered epithelial-stromal interactions specific to deep infiltrating disease (Crean and Murphy, 2021).

Our study findings are consistent with the prevailing notion that bidirectional interactions between endometrial-type epithelium/stroma and the local microenvironment play critical roles in endometriosis pathogenesis. We discovered context-specific features of mesothelial cells, which were less differentiated when associated with endometrioma and some peritoneal lesions. A set of atypical mesothelial cells that co-express mesothelial markers (*WT1* and *PDPN*), hormone receptors (*ESR1* and *PGR*), *KRT10* and *ACTA2* were enriched in endometrioma specimens. These altered mesothelial cells may contribute to adhesion formation, for example due to downregulated expression of the cell surface glycoprotein mesothelin. Arguably the greatest insight into microenvironmental remodeling was revealed when we interrogated the transcriptional hallmarks of *ARID1A* mutant and wild-type endometrial-type epithelium. We observed dysregulated gene expression in endothelial associated with *ARID1A*-mutant epithelium, likely mediated by SOX17, a transcription factor which when expressed in Müllerian epithelium, induces a pro-angiogenic gene signature and altered secretion of angiogenesis-regulating proteins (Chaves-Moreira et al., 2020). Further investigations will be needed to functionally validate the impact of *ARID1A* and *KRAS* mutations on the behavior of endometriosis epithelial cells and altered interactions with microenvironmental populations, and to understand the specific context in which *ARID1A* mutations, in particular, result in endometriosis-associated ovarian cancer (Anglesio et al., 2015; Wiegand et al., 2010).

There are caveats to this study. With an overall cohort size of nine endometriosis patients and two unaffected women we were underpowered to test for confounding effects of age or identify associations between molecular features and patient symptoms or outcomes. Representative ‘normal’ tissue from women with and without endometriosis is challenging to obtain and therefore underrepresented in this study, particularly uninvolved peritoneum and pre-menopausal ovary tissue. We attempted to profile uninvolved peritoneum from endometriosis patients by including four samples from two endometriosis patients where no endometriosis was detected upon pathologic review, however in these cases we saw some evidence of endometrial-type epithelium and endometrial-type stroma, suggesting the portion of the tissue used for scRNA-seq did indeed contain endometriosis tissue. While scRNA-seq is unlikely to have utility as a diagnostic tool, this illustrates the challenge of obtaining a pathologic confirmation of endometriosis, particularly in cases with only a few small lesions suspected at laparoscopy that may be ‘missed’ upon pathologic review due to intrinsic limitations of the embedding and sectioning processes.

Endometriosis research has been substantially hindered by challenges in generating global molecular profiles of tissues. This single cell atlas of endometriosis therefore represents a valuable and timely resource for the endometriosis research community. Continued large-scale somatic profiling efforts are clearly warranted, as these data indicate that endometriosis likely comprises multiple subtypes that will likely require different approaches to treatment and diagnosis.

## Methods

| **Reagent or Resource** | **Source** | **Identifier** |
| --- | --- | --- |
| **Antibodies** |  |  |
| Anti-SOX17 | R&D Systems | AF1924; RRID: AB_355060 |
| Anti-CAPS | Sigma-Aldrich | HPA043520; RRID:AB_10964138 |
| Anti-ARID1A | Abcam | EPR13501 (ab182560) |
| **Biological Specimens** |  |  |
| Human endometriosis, endometrium and ovary tissues (fresh) | Cedars-Sinai Medical Center – Divisions of Minimally Invasive Gynecologic Surgery and Gynecologic Oncology | This study |
| Human endometriosis and endometrium (formalin fixed) | Cedars-Sinai Medical Center – Divisions of Minimally Invasive Gynecologic Surgery and Gynecologic Oncology and Department of Pathology and Laboratory Medicine | This study |
| **Chemicals, peptides, and recombinant proteins** |  |  |
| 10X Collagenase/Hyaluronidase in DMEM | STEMCELL Technologies | 07912 |
| Minimal Essential Medium (MEM) | Corning | 10-010-CV |
| Deoxyribonuclease I | Sigma-Aldrich | DN25-1G |
| **Critical commercial assays** |  |  |
| Dead Cell Removal Kit | Milteny Biotec | 130-090-101 |
| Chromium Single Cell 3' GEM, Library & Gel Bead Kit v3 | 10x Genomics | 1000075 |
| Chromium Single Cell A Chip Kits | 10x Genomics | 120236 |
| Chromium Chip B Single Cell Kit | 10x Genomics | 1000153 |
| **Software and algorithms** |  |  |
| Cell Ranger version 3.1.0 | https://support.10xgenomics.com/single-cell-gene-expression/software/overview/welcome | RRID:SCR_017344 |
| R package Seurat version 3.2.2 | https://satijalab.org/seurat/ | RRID:SCR_007322 |
| R package DoubletFinder | https://github.com/chris-mcginnis-ucsf/DoubletFinder | RRID:SCR_018771 |
| R package MAST | https://github.com/RGLab/MAST | RRID:SCR_016340 |
| R package Monocle3 | https://cole-trapnell-lab.github.io/monocle3/docs/introduction/ | RRID:SCR_018685 |
| R package SCENIC version 1.2.0 | https://rawcdn.githack.com/aertslab/SCENIC/6aed5ef0b0386a87982ba4cc7aa13db0444263a6/inst/doc/SCENIC_Running.html | RRID:SCR_017247 |
| R package - MUSIC | https://github.com/xuranw/MuSiC/ | RRID: SCR_008792 |
| R package ReactomePA | https://bioconductor.org/packages/release/bioc/html/ReactomePA.html | N/A |
| R package Harmony | https://portals.broadinstitute.org/harmony/articles/quickstart.html | N/A |

### Resource availability

#### Lead contact

Further information and requests for resources and reagents should be directed to and will be fulfilled by the Lead Contact, Kate Lawrenson.

#### Materials availability

This study did not generate new unique reagents.

#### Data and code availability

The data generated during this study are available at NCBI GEO under the following accession number: XXX (to be provided by time of publication).

### Patient specimens

This project was performed with approval of the Institutional Review Board at Cedars-Sinai Medical Center. All patients provided informed consent. Human endometriosis, endometrial or ovarian tissues were placed in sterile serum-free MEM at 4°C and transferred to the tissue culture laboratory.

### Method Details

#### Surgical and pathologic review

All patients presented to our minimally invasive gynecologic surgery division at Cedars-Sinai Medical Center for consultation. The division comprises three fellowship-trained minimally invasive gynecologic surgeons who perform over 150 advanced endometriosis procedures annually. Patients were evaluated by the surgeon with a detailed history and physical taken as well as all imaging reviewed, and the decision was made for surgery. Surgery was performed either laparoscopically or robotically, with identification and excision of any obvious or suspected lesions. Lesions were excised using either ultrasonic energy or monopolar energy. Wide margins were attempted for each excision. For example, a lesion in the ovarian fossa would lead to the full peritoneum in the ovarian fossa being removed. Deep infiltrating endometriosis resections were performed until normal anatomy was restored, leaving the endometriosis lesion intact. Ureterolysis and mobilization of the rectosigmoid colon were performed when necessary. Areas were separately labeled and passed off to nursing for the research study.

#### Tissue collection

A collection protocol was created and implemented in the pathology laboratory to ensure collection of endometriosis samples from consented patients without risk of compromising the ability to provide clinical histopathologic diagnoses on resected tissue (Figure S6). Examination of 5 deeper sections cut at 50 μm intervals was performed in all cases in which endometriosis was not identified in the first level.

#### Tissue processing

Tissues were minced into ~1-2 mm pieces and digested with 1× Collagenase/Hyaluronidase (STEMCELL Technologies) and 100 μg/mL DNase I (Sigma-Aldrich) in 7mL serum-free MEM. The sample was incubated at 37°C with constant rotation for 90 mins. The supernatant was harvested, and the cell suspension was spun at 300 g for 10 mins at 4°C. To lyse red blood cells, the cell pellet was resuspended in an RBC lysis buffer (0.8% NH_4_Cl, 0.1% KHCO_3_, pH=7.2) and incubated for 10 mins at room temperature. Cell suspensions were spun again at 300 g for 10 mins at 4°C and the cell pellet was resuspended in phosphate buffered saline (PBS), or, if >5% dead cells were observed by trypan blue staining, cells were resuspended in dead cell removal buffer and dead cell removal was performed according to manufacturer’s instructions.

Remaining cells were frozen in 90% fetal bovine serum with 10% DMSO in a Mr. Frosty container placed at −80°C. Frozen cell vials were stored in LN_2_. Cells were thawed and transferred into a conical tube with 7 mL serum free media and spun at 300 g for 10 mins at 4°C. The cell pellet was resuspended in 100 μL PBS. Cells were counted using a hemocytometer and the sample volume adjusted to achieve a cell concentration between 100/μL to 2,000/μL.

#### Single cell capture, library preparation and next-generation sequencing

Single cells are captured and barcoded using 10X Chromium platform (10X Genomics). scRNA-seq libraries were prepared following the instructions from Chromium Single Cell 3’ Reagent Kits User Guide (v2 or v3). Briefly, Gel Bead-In EMulsions (GEMs) are generated using single cell preparations. After GEM-RT and cleanup, the cDNA from barcoded single cell RNAs were amplified before quantification using Agilent Bioanalyzer High Sensitivity DNA chips. The single cell 3’ gene expression libraries were constructed and cDNA corresponding to an insertion size of around 350 bp selected. Libraries were quantified using Agilent Bioanalyzer High Sensitivity DNA chip and pooled together to get similar numbers of reads from each single cell before sequencing on a NovaSeq S4 lane (Novogene).

#### Single cell data processing and filtering

Raw reads were aligned to the hg38 reference genome, UMI (unique molecular identifier) counting was performed using Cell Ranger v.3.1.0 (10X Genomics) pipeline with default parameters. Normal ovary from patient 3 was removed due to a low fraction of reads in cells (sample proportion: 35.3%, ideal fraction >70%). Left pelvic side wall from patient 10 was removed due to a low fraction of reads mapped to the transcriptome (sample proportion: 17.0%, ideal fraction >30%) and a low fraction of reads in cells (sample proportion: 38.7%, ideal fraction >70%). For each individual sample we removed cells with high mitochondrial content (>20%) and cells with less than 200 genes. We applied DoubletFinder (McGinnis et al., 2019) following the expected percentage of doublets for each sample (0.8% per 1000 cells) to remove potential doublets based on the expression proximity of each cell to artificial doublets. Combined Seurat objects were adjusted for bias aiming to remove confounding factors that are potential sources of variation. We considered: sequencing batch, number of reads, mitochondrial mapping percentage and sample as parameters for Seurat *SCTransform* function. We applied the *CellCycleScoring* Seurat procedure to check if genes related to cell cycle are guiding the PCs and found that none of the 20 first PCs had cell cycle genes within the top 20 positive and negative genes. To integrate the 30 samples we used Harmony (Korsunsky et al., 2019), with *lambda = 0.2*, to reduce technical batch effects. We then used the reduced Seurat object to define an initial cluster considering Seurat’s *FindNeighbors* (using 25 dimensions as parameter) and *FindClusters* function with a resolution of 3.

#### Identification of major cell types and epithelial subgroups

To define the major cell type for each cluster we divided our procedures in two steps. On step one we performed differential expression analysis (one *versus* all) using MAST (Finak et al., 2015), implemented in the FindAllMarker and FindMarker functions in Seurat. Then, we checked the presence of the following marker genes for global annotation of cell types that were differentially expressed at log_2_ fold change 0.2 and adjusted p-value 0.05. Epithelial cells (*EPCAM*, *KRT8*, *KRT18*, *KRT19*, *KRT7*, *KRT10*), Fibroblasts (*DCN*, *COL11A2*, *FAP*, *PDGFRA*, *COL11A1*, *COL1A1*, *PDGFRB*), Myeloid cells (*LYZ*, *CD14*, *MME*, *C1QA*, *CLEC10A*), Endothelial cells (*CLDN5*, *PECAM1*, *CD34*, *ESAM*), Plasma cells (*JCHAIN* plus *CD79A*), B cells (*JCHAIN*), Smooth muscle cells (*ACTA2*), Mast (*TPSB2*), Erythrocytes (*HBB*, *GYPA*), T cells (*CD2*, *CD3D*, *CD3E*, *CD3G*, *CD8A*, *CCL5*) and Natural Killer cells (*TYROBP*, *FCGR3A*). For each cluster we build a matrix of differentially expressed gene counts by normalizing each count with the total markers in each cell type (Figure S2). To assign the cell type based on the matrix of DEG counts we applied the following rules. First, clusters that only had cell type specific genes for one cell type contained within the DEG list were assigned to the corresponding cell type. If the cluster *i* has >35% of the cells expressing at least one keratin gene and the average of scaled expression is greater than 1 then the Epithelial cell type was assigned to cluster *i*. If the cluster *i* has multiple cell type markers contained within the DEG list, we first checked if *ACTA2* was expressed, if so we checked if the proportion of Fibroblast markers was greater than 25%, then assigned the cluster *i* as Fibroblast otherwise as Smooth muscle cells. For the case where we have multiple markers but no *ACTA2* we then checked which marker has max proportion and assigned the correspondent cell type to cluster *i*, if the multiple markers had the same proportion we skipped the assignment for cluster *i*. Clusters with no markers counts were also not assigned a cell type with this decision tree. The next step aimed to identify cell types for the clusters that did not express canonical patterns of expression of known cell-type specific genes. We use the 84 clusters for which we could successfully assign a cell type in step one as a reference panel. We selected up to 100 of the top-ranked DEG (log_2_ FC > 0; p <0.05) for each cluster to calculate pairwise Pearsons’ correlations across all clusters (to create a union set of 1,960 genes). Clusters with no cell markers were assigned the identity associated with the most correlated cell type with known identity. Cellular transcriptomes of cells in C1 correlated with cells in C0 - Fibroblasts (r=0.91, Pearson correlation), C12 correlated with 41 - Epithelial cells (0.97), C29 correlated with C0 - Fibroblasts (0.89), C37 correlated with C65 - Epithelial cells (0.96), C41 correlated with C12 - Epithelial cells (0.97), C55 correlated with C50 - Smooth muscle cells (0.8), C63 correlated with C7 - Epithelial cells (0.8), C65 correlated with C37 - Epithelial cells (0.96), C83 correlated with C37 - Epithelial cells (0.92), C92 correlated with C37 Epithelial cells (0.88), C95 correlated with C83 - Epithelial cells (0.77), C84 20% of cells expressed *KRT10* and it is correlated with C88 - Epithelial cells (0.82), C4: 23% cells expressed *KRT10* and it is correlated with C88 - Epithelial cells (0.81) C12 21% cells expressed *KRT10* and it is correlated with C88 - Epithelial cells (0.77) (Figure S2). UMAP (uniform manifold approximation and projection)(Becht et al., 2018) was used for visualizing cell types and clusters with representative markers.

#### Epithelial and fibroblast subgroup analyses

Epithelial and fibroblast clusters were identified from the parent cluster and analysed in isolation. We defined the cell clusters considering Seurat’s *FindNeighbors* (using 20 dimensions as parameter) and *FindClusters* function with a resolution of 0.5. The eutopic endometrium sample from patient 11 exhibited a markedly different epithelial profile that dominated many of the subclusters when included, and so this sample was removed from the epithelial-specific analyses. Pathway analyses were performed using Reactome (Yu and He, 2016).

#### Transcription factor analyses using SCENIC

Key transcription factors and their gene regulatory network (GRN) were identified using SCENIC R package v.1.2.0 (Single Cell rEgulatory Network Inference and Clustering (Aibar et al., 2017)) considering hg38 reference genome from RcisTarget database. We applied SCENIC to endometrial-type epithelium and stroma using the normalized expression matrix from Seurat as the input matrix.

#### Immunohistochemistry for ARID1A, CAPS and SOX17

Immunohistochemistry assays for ARID1A were performed on 5μm tissue sections on Superfrostplus slides and used as surrogate for somatic loss-of-function alterations following established standards for staining and scoring (Khalique et al., 2018). ARID1A staining was performed on a Leica Bond Rx (Leica Biosystems) using rabbit monoclonal antibody EPR13501 (Abcam) at 1:3000 dilution. Slides were scored by pathologist A.E-N, assessed for (loss of) nuclear staining in epithelium with retained stromal nuclear staining serving as an obligate internal control. CAPS staining was performed using HPA043520 (Sigma Aldrich) at a 1:5,000 dilution and SOX17 staining performed using goat polyclonal antibody AF1924 (R&D Systems). SOX17 and CAPS staining was performed on a Ventana Discovery Ultra autostainer (Roche Ventana). CAPS and SOX17 stained slides were scored by pathologist F.M. For SOX17 the proportion of positively stained epithelium was estimated, and the predominant straining intensity categorized as negative, weak, moderate or strong.

#### KRAS mutation testing by ddPCR

Endometrial glands and stroma were enriched from 10% dilute H&E stained, 5-7μm sections by needle macrodissection with a 20-gauge needle. DNA was then extracted using the Arcturus PicoPure DNA Extraction Kit (Thermo) and quantified using the Qubit 2.0 Fluorometer (Thermo). 2ng of DNA was pre-amplified from the KRAS G12 codon region in a 20uL reaction vol with TaqMan Genotyping Mastermix (Thermo); forward and reverse KRAS primers (IDT; Table S16). The following pre-amplification conditions were used: 95°C - 10 min; 10 cycles of 94°C/30s, 60°C/4min on an AC4 thermal cycler (FroggaBio). 5-fold diluted pre-amplified DNA was used in multiplex ddPCR and subsequently individual-variant ddPCR for validation (if positive). All ddPCR reactions used the same flanking primers (Table S16) and ddPCR Supermix for Probes (no dUTP; Biorad) in a 25μl reaction vol with cycling parameters 95°C/10 min, followed by 40 cycles of 94°C/30s, 60°C/90s, on an AC4 thermal cycler (FroggaBio). Droplets were generated on the BioRad QX200 Automated Droplet Generator and read on the Biorad QX200 Droplet Reader. Multiplex ddPCR included an equimolar mix of probes for KRAS G12C/D/R and was used for detection of KRAS G12C/D/R/V/A/S based on counts and cluster position of fluorescence signal. Any positive multiplex assay was then validated with individual variant ddPCR reactions (see Table S16 for probes).

The following limits of detection thresholds were applied in the multiplex assay: variant allele frequency (VAF) threshold for KRAS G12C/D/R/A/S allele needed to be 3x average of negative control reactions for the given allele. For KRAS G12V allele (VAF) was required to be 1x average of negative controls. For individual allele variants, ddPCR 3x average of negative control reactions was used universally as the minimum detection threshold.

#### Defining pseudotime cell trajectories using Monocle 3

We used Monocle 3 (Cao et al., 2019; Qiu et al., 2017; Trapnell et al., 2014), implemented in R, to infer cell trajectories. We extract counts, phenotype data, and feature data from the Seurat object after filtering the cells of interest. To normalize and pre-process the data we considered *num_dim = 100*. We cluster the cells using Monocle procedures and applied the *learn_graph* procedure using *use_partition* as FALSE to keep a unique trajectory. To define “roots” of the trajectory we adapted the helper function, available on Monocle 3 website, to identify the root principal points by informing a cluster of interest.

#### Deconvolution analysis using MuSiC

To compare the profile of bulk tumor tissues and the histologic components present in each specimen of this study, we used the multi-subject single-cell deconvolution - MuSiC method (Wang et al., 2019). First, we selected all epithelial cells present in Endometrioma, Extra-ovarian Endometriosis and Eutopic Endometrium specimes. The gene signature was based on the common genes expressed on both scRNA-seq and bulk RNA datasets. We downloaded data from the following GEO datasets: GSE129617 (n=25): 24 primary tumors and 1 ascites samples; GSE73614 (n=107): 24 CCOC and 35 Endometrioid; GSE9899 (n=285): from 243 Ovary samples we selected 20 endometrioid samples.

## Acknowledgements

Some of the specimens were collected as part of the Biologic and Epidemiologic Markers of Endometriosis (BEME) study. We wholeheartedly thank all the patients who donated the specimens used in this study, and the team at the Biobank and Translational Research Laboratory who supported tissue procurement and histologic analyses. We thank Christine Chow and Monica Ta at the Genetic Pathology Evaluation Centre (GPEC) for technical support for the mutation analyses.

